# A 3D Topographical Model of Parenchymal Infiltration and Perivascular Invasion in Glioblastoma

**DOI:** 10.1101/298240

**Authors:** Kayla J. Wolf, Stacey Lee, Sanjay Kumar

**Author notes:** Corresponding author. *Email address:* (S. Kumar).

## Abstract

Glioblastoma (GBM) is the most common and invasive primary brain cancer. GBM tumors are characterized by diffuse infiltration, with tumor cells invading slowly through the hyaluronic acid (HA)-rich parenchyma toward vascular beds and then migrating rapidly along microvasculature. Progress in understanding local infiltration, vascular homing, and perivascular invasion is limited by an absence of culture models that recapitulate these hallmark processes. Here we introduce a platform for GBM invasion consisting of a tumor-like cell reservoir and a parallel open channel “vessel” embedded in 3D HA-RGD matrix. We show that this simple paradigm is sufficient to capture multi-step invasion and transitions in cell morphology and speed reminiscent of those seen in GBM. Specifically, seeded tumor cells grow into multicellular masses that expand and invade the surrounding HA-RGD matrices while extending long (10-100 µm), thin protrusions resembling those observed for GBM *in vivo*. Upon encountering the channel, cells orient along the channel wall, adopt a 2D-like morphology, and migrate rapidly along the channel. Structured illumination microscopy reveals distinct cytoskeletal architectures for cells invading through the HA matrix versus those migrating along the vascular channel. Substitution of collagen I in place of HA-RGD supports the same sequence of events but with faster local invasion and a more mesenchymal morphology. These results indicate that topographical effects are generalizable across matrix formulations, but that mechanisms underlying invasion are matrix-dependent. We anticipate that our reductionist paradigm should speed the development of mechanistic hypotheses that could be tested in more complex tumor models.

## Introduction

Glioblastoma (GBM) is the most common and malignant primary brain tumor, with a median survival time of 14 months and comparatively little improvement in clinical outcome over the past few decades.^1^ GBM is characterized by a diffuse, infiltrative pattern of spread in which tumor cells evade surgical resection by migrating away from the tumor mass and resist chemotherapy and radiation.^2,3^ Cell interactions with extracellular matrix (ECM) represent a relatively unexplored potential therapeutic target. Tumor cells hijack ECM to promote survival and invasion, and disruption of cell-ECM interactions shows promise for sensitizing cells to therapeutic intervention.^4^ An important detail of brain ECM is that its composition varies dramatically by region. The ECM surrounding microvasculature is rich in collagen, fibronectin, and laminin, while the parenchymal space is generally devoid of these proteins and instead rich in hyaluronic acid (HA), which both supports direct cell adhesion and organizes other matrix components such as tenascin.^5–7^ Tumor cells invade slowly through the hyaluronic-acid rich parenchyma and then rapidly along vascular tracks, analogous to cars moving on a highway. ^8–12^ Chemotactic signals from endothelial and other stromal cells and haptotactic signals from the vascular ECM comprise a perivascular niche (PVN) that promotes invasion and resistance to therapy. ^5,13–17^

Despite the established role of the PVN in driving rapid dissemination, mechanisms governing the transition from intraparenchymal to perivascular invasion are not well understood. Investigation of local infiltration, vascular homing, and perivascular invasion is made challenging by the absence of advanced culture models.^18–20^ Most studies are performed *in vitro* using rigid, 2D culture dishes. While these simplified paradigms have high scalability and throughput, they do not allow one to control functionally important properties of the brain microenvironment such as stiffness, microarchitecture, dimensionality and microregional heterogeneity. For example, GBM cells *in vivo* form microtubes that can signal through interpenetrating networks which are not observed in 2D culture.^21^ Animal models offer a fully integrated system but are not amenable to parallelized discovery and screening and lack the tunability needed to quantitatively dissect invasion mechanisms.^20,22,23^

3D culture models derived from natural or synthetic ECM components offer a compromise, in that these systems capture structural aspects of matrix relevant to invasion while retaining some tunability and scalability. For example, implantation of tumorspheres into 3D hydrogels enables integrated investigation of how dimensionality, stiffness, and microarchitecture control invasion.^24^ With these concepts in mind, our laboratory has explored the utility of 3D HA hydrogels for investigating GBM invasion. The nanoporous microarchitecture of HA is reminiscent of brain parenchyma, and the elastic modulus is similar to that of brain tissue (300 – 3000 Pa).^25,26^ Moreover, morphologic hallmarks of GBM intraparenchymal invasion seen in brain can be recapitulated in 3D HA-RGD gels.^27,28^ Still, 3D tumorsphere assays are spatially uniform and thus do not capture the structurally heterogeneous tracks that are closely associated with invasion in vivo.

Early efforts to build microstructural cues into culture models of tumors including GBM have shown great promise in elucidating mechanisms of invasion. For example, confined microchannels have been used to investigate regulation of nuclear squeezing during 3D invasion.^29–33^ The heterogeneity between the vascular basement membrane and parenchyma has been modeled by layering matrix types, revealing distinct differences in migration depending on the composition of each layer.^34,35^ Cells also rapidly follow anatomical tracks, modeled by micropatterned adhesive ligands and electrospun fibers.^36,37^ For perivascular invasion specifically, microfluidic devices have been developed to investigate vascular homing and extravasation using separate chambers for endothelial cells, 3D matrix, and a cell reservoir.^18,38–42^ The devices do not incorporate the cylindrical geometry of vasculature embedded within 3D matrix, which is vital to understanding how GBM interacts with anatomical tracks. Furthermore, the 3D matrices applied in these systems are often based on fibrillar collagens, which tend not to be abundant in brain outside of vascular compartments. Finally, the chambers are not typically embedded in matrix, which in principle could allow some cells to invade along the matrix-wall interface instead of invading in 3D until reaching the matrix-vasculature interface. A model of invasion, therefore, should include both a 3D HA-rich matrix to capture aspects of vascular homing and topographical cues to investigate migration along anatomical tracks.

Here, we develop a simple 3D topographical model that enables us to recapitulate multiple stages and features of GBM progression during vascular homing and subsequent migration. Specifically, we find that the incorporation of an open channel as a vessel mimic parallel to a cell reservoir enables imaging of mass expansion, slow invasion through 3D matrix, and rapid invasion along the channel wall. In a proof-of-principle demonstration, we show that arrival at the vascular channel is accompanied by a transition in tumor cell morphology and invasion speed that is broadly reminiscent of perivascular homing and invasion in GBM. Furthermore, we find that while the overall cell speeds and actin cytoskeletal morphologies underlying the transition are dependent on matrix type, the relative transition in speed is generalizable across matrices. Thus, while matrix formulation influences mechanisms of invasion, topography can influence invasion within a particular matrix type.

## Methods

### HA Matrix Synthesis

HA hydrogels were synthesized as previously described.^27^ Briefly, methacrylic anhydride (Sigma-Aldrich, 94%) was used to functionalize sodium hyaluronate (Lifecore Biomedical, Research Grade, 66 kDa - 99 kDa) with methacrylate groups (Me-HA). The extent of methacrylation per disaccharide was quantified by ^1^H NMR as detailed previously^43^ and found to be ~85% for materials used in this study. To add integrin-adhesive functionality, Me-HA was conjugated via Michael Addition with the cysteine-containing RGD peptide Ac-GCGYGRGDSPG-NH2 (Anaspec) at a concentration of 0.5 mmol/L. Finally, 3wt% Me-HA was crosslinked in phenol-free DMEM (Invitrogen) with bifunctional thiol dithiothretiol (DTT, Sigma-Aldrich). A concentration of 19 mmol/L DTT was selected to yield a shear modulus of ~300 Pa. After 1 hr of crosslinking, the hydrogels were rinsed and soaked in room temperature PBS for 1 hr before cell seeding.

### Rheology Characterization

The shear modulus of hydrogel formulations was measured using oscillatory rheometry (Anton Parr Physica MCR 310) as described previously.^27^ Briefly, hydrogels were first crosslinked by incubation for 1 hr in a humidified 37˚C chamber. Rheological testing consisted of frequency sweeps ranging from 100 to 0.1 Hz at 0.5% amplitude also in a humidified 37˚C chamber. Shear modulus was reported as the average storage modulus for 3 tests per type of matrix composition at an oscillation frequency of 0.5 Hz.

### Collagen Matrix Synthesis

Rat tail collagen I (BD Biosciences) was used to form hydrogels according to manufacturer’s instructions. Briefly, a solution of 1v/v% 1 N NaOH (Carolina Biological Supply), 10v/v% 10x PBS (Fisher BioReagent), and 50 v/v% of 3.84 mg/mL cold collagen I in sterile distilled water was mixed thoroughly on ice. The solution was then pipetted into the desired mold and incubated for 30 minutes at 37 °C. Finally, solutions were rinsed and soaked for 1 hr in room temperature PBS before cell seeding.

### Cell culture

U87-MG human glioblastoma cells were obtained from the University of California, Berkeley Tissue Culture Facility, which sources its cultures directly from the ATCC. Cells were cultured in DMEM (Invitrogen) supplemented with 10% calf serum (JR Scientific), 1% penicillin-streptomycin, MEM nonessential amino acids, and sodium pyruvate (Invitrogen). Cells were harvested using 0.25% trypsin-EDTA (Thermo Fisher Scientific). Cells were screened bimonthly for mycoplasma and authenticated annually via short tandem repeat analysis. For invasion in PVN model devices, 0.2 µL of cells at 5×10^6^ cells/mL were injected. Devices were cultured in 6-well plates with medium changes every 3-4 days. In devices treated with blebbistatin, medium containing 10 µM blebbistatin was added to the entire well at day 1 and exchanged every 3 days. In other devices, recombinant human epithelial growth factor (EGF) (R&D Systems) at 2 µg/mL concentration in 10% serum-containing medium was added to the vessel-like channel and exchanged every 3 days.

### Device Fabrication

First, the PDMS support was fabricated by mixing a 10:1 mass to mass ratio of Sylgard 184 elastomer with initiator (Dow Corning). The mixture was pipetted into the desired mold and cured at 80 °C for 2 hr. Horizontal PDMS spacers were first created to separate wires (167 µm diameter, Hamilton) by 500 µm using 500 µm outer diameter glass capillaries (CTech Glass) without attention to height in the *z* direction, and then sliced and used to space the wires for all subsequent fabrication. The PDMS supports for hydrogels were fabricated by aligning wires with pre-fabricated horizontal spacers to separate the parallel wires at each end. The wires were spaced between glass coverslips and suspended using 170 µm PDMS strips as vertical spacers. After curing, a 3 mm hole punch and razor blade was used to cut the center of the PDMS support and create space for the hydrogel. The PDMS support was assembled between two 18 mm #1 glass coverslips (VWR) and fastened with a drop of 5-minute epoxy (ITW Devcon) at two corners.

To cast the gel, the wires were first reinserted into the assembled device to create a mold. Next, the hydrogel matrix was inserted in the side of the device. Wires were removed after the hydrogel solution had solidified leaving two open channels. After rinsing and soaking the hydrogels, cells were inserted into the freshly fabricated device using a syringe (Hamilton). Cut wires were used to plug each end of the cell reservoir, and the entire device was placed into the bottom of a 6-well plate and bathed in 3 mL of medium. Cells and gels were equilibrated in medium overnight before imaging. Medium was changed every 3-4 days. To introduce diffusible soluble factors into the vessel-like channel, a 33-gauge syringe needle (Jensen Global) pre-loaded with growth factor and medium of interest and was then inserted into the PDMS support. The opposite end of the open channel was plugged with an additional wire. Diffusion between the open channel and the surrounding bath of medium was blocked in some devices by inserting wires into the PDMS support, fully occluding the channel.

### Invasion Analysis

For area analysis in HA PVN devices, cells in devices were imaged once every 3 days using Eclipse TE2000 Nikon Microscope with a Plan Fluor Ph1 10x objective. Images were acquired and large images were stitched using NIS-Elements Software. For each device, the cell reservoir and area of invasion were outlined in ImageJ and normalized to the total cell area from day 1, assumed to be reservoir only. Migration assays were performed by imaging at 15 minute intervals for 6 hrs. The ImageJ plugin Manual Tracking was used to track cell movements in each frame and calculate an average cell speed. To analyze single-cell speed transitions, cells were tracked for at least 6 hr until a transition event was observed. Only the 3 hr prior to and directly after a transition were analyzed. The transition point was defined as the time in which the entire nucleus had exited the 3D matrix. Average speed was calculated for cells before and after the transition. All cell motility imaging was performed at least 1 day after seeding in collagen gels, and at least 10 days after seeding in HA-RGD gels.

### Cell labeling and Confocal Fluorescence Microscopy

Cells in 3D matrices were fixed with a 4 w/v% paraformaldehyde solution (PFA, Alfa Aesar) for 4 hr and then permeabilized with 0.5 v/v% Triton-X (Thomas Scientific) solution for 1 hr. Cells were labeled with 1 µg/mL 4′,6-diamidino-2-phenylindole (DAPI, Sigma-Adrich) and 488-labeled phalloidin (Cytoskeleton) by soaking overnight and then washed with PBS thoroughly overnight. After fixation and immunostaining, confocal images were acquired on a Zeiss LSM 780 NLO Axioexaminer upright microscope equipped with a 32-channel GaAsP detector and 2 PMTs using a PlanApo Chromat 20x/1.0 water-dipping objective. Samples were illuminated using an Argon multiline laser for excitation at 488nm and a 405nm diode laser. Zen 2010 acquisition software was used. Finally, z slices separated by 5 µm were projected in ImageJ using 3D volume viewer. For investigation of cell division within the tumor reservoir, devices that had been cultured for >16 days were disassembled and fixed with 4 w/v% PFA in PBS. The HA matrix was treated with 2500 U/mL hyaluronidase from bovine testes (Sigma) overnight to improve antibody diffusion into the fixed sample. Cells were then labeled with 1 µg/mL 4′,6-diamidino-2-phenylindole (DAPI, Sigma-Adrich), Alexa Fluor 546-phalloidin (Thermo-Fisher), monoclonal rabbit anti-Ki-67 primary antibody (Abcam, clone SP6), and Alexa Fluor-647 polyclonal goat anti-rabbit secondary antibody (Abcam). Stacked confocal images of Ki-67 labeled cells were obtained with a swept-field confocal microscope (Prairie Technologies) with an Olympus LUM Plan FL 60x water immersion objective.

### Live/Dead Cell Viability Assay

To perform a cell viability assay (Invitrogen), devices that had been cultured for >16 days were disassembled, and cells in 3D matrices were washed with PBS at 37 ºC for 10 minutes. Gels with cells were then incubated in 2 µM calcein AM and 1 µM ethidium homodimer-1 in PBS for 20 minutes with gentle rocking. Finally, gels were rinsed for 5 minutes in PBS prior to imaging using an Eclipse TE2000 Nikon Microscope with a Plan Fluor Ph1 10x objective.

### Structured Illumination Microscopy (SIM)

For SIM imaging, devices were disassembled by removing the top coverslip and placing the sample face-down onto a #0 coverslip dish (MatTek). To image cells within channels, gels were trimmed with a scalpel and then placed into the dish with PBS for imaging. Cells were imaged directly using an Elyra structured illumination microscope (Zeiss) and a Plan-Apochromat 63x/1.4 Oil DIC M27 objective (Zeiss). Samples were illuminated using an Argon multiline laser for excitation at 488nm and a 405nm diode laser. Zen 2010 software was used for image acquisition. For z-stack images, slices were 1 µm apart. Finally, z-projections were formed in ImageJ with 2×2 binning using 3D project. Artifacts caused by imaging deep with 3D HA were minimized by filtering the Fourier transform for images of cells within the HA channel.

### Statistical analysis

The sample numbers necessary to obtain a power of 0.8 were estimated prior to experimentation based on estimates from previous studies (G* Power).^44^ Each device was fabricated and seeded independently. GraphPad Prism 7 software was used to graph data and perform statistical analysis. Reservoir and invasion areas in each device over time were compared using the Friedman test followed by Dunn’s multiple comparison test. Bias in area of invasion was compared using a paired t-test. Migration speeds within the matrix and channel were compared as the average of 5 cells in 3 independent devices for each condition to give n=15 where variability is primarily from cell to cell. Several exclusion criteria were applied when selecting cells for analysis: cells must be clearly visible in all frames of imaging; cells must be leading invasion in the 3D matrix or actively moving in the open channel; and cells must not undergo division during the imaging interval. Data were compared using a student’s t-test. For single cell transitions, 2-3 cells were measured in 3 independent devices to give a total of n=7. Data were compared using a paired t-test.

### Approvals

This study did not involve human subjects research, and so no ethics approval was required.

## Results and Discussion

### Mimicking Vascular Tracks Embedded within an HA Matrix with a Simple Device

GBM invasion is characterized by tumor expansion, slow invasion through the HA-rich parenchyma, and rapid invasion along vascular tracks **(Fig 1A).** During this process, cells transition from 3D migration mode to a 2D-like migration mode as they follow the interface between the basement membrane and surrounding parenchyma.^8,11,35^ We hypothesized that a simple topographic model of the PVN composed of a cell reservoir parallel to an open channel would induce similar progression in invasion **(Fig 1B)**. Specifically, we hypothesized that by restricting cell invasion to the 3D matrix until cells encountered the open channel, cells would transition from slow migration through the 3D matrix to more rapid migration along the 2D wall of the open channel, analogous to GBM invasion kinetics in vivo.

**Figure 1.**
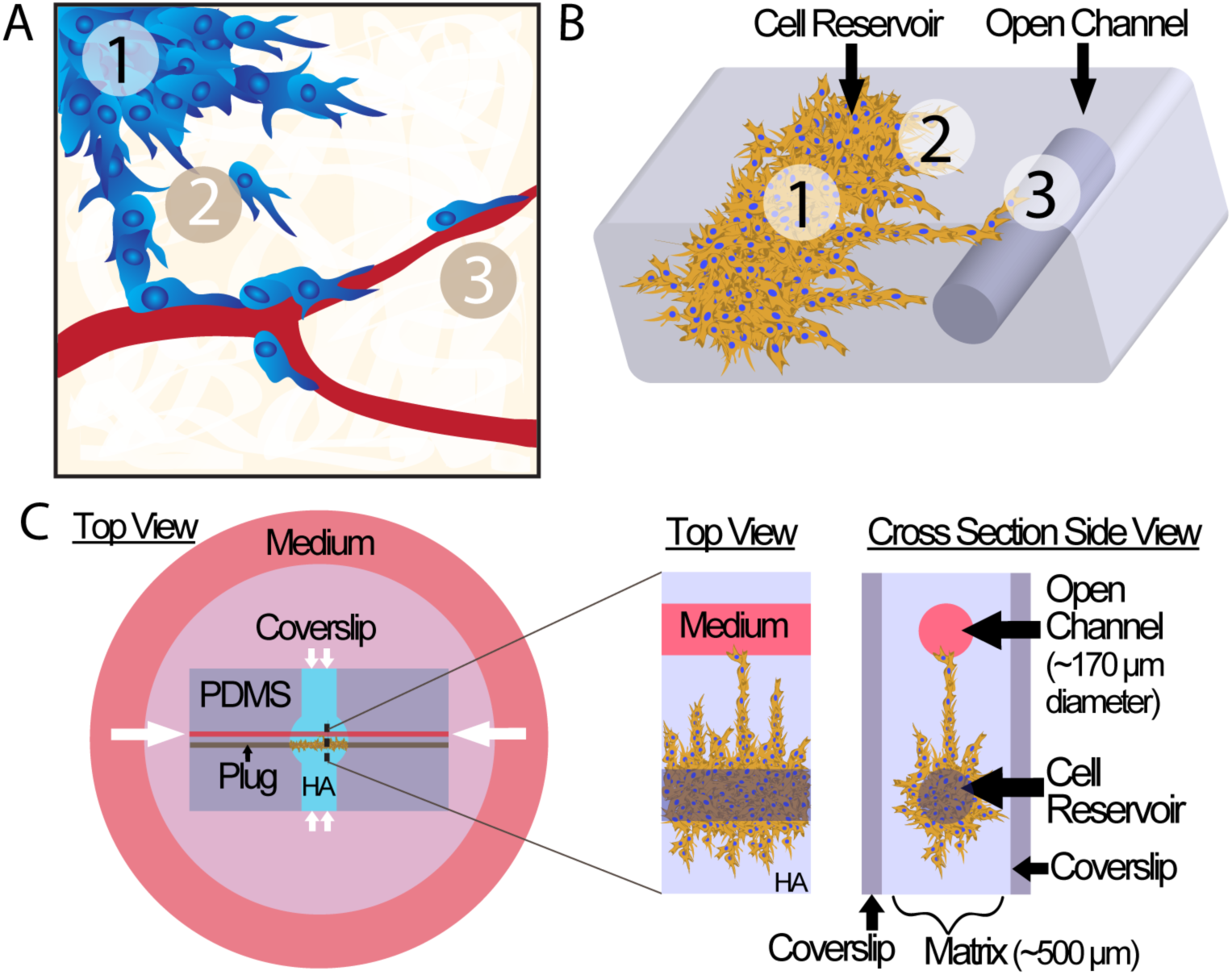
A topographical model of the perivascular niche. A) Schematic of GBM progression representing 1) tumor expansion, 2) slow invasion through the HA-rich parenchyma, and 3) rapid migration along vascular tracks. B) GBM progression can be modeled using a simple 3D topographical model of a vessel. C) Device Schematic. Hydrogel matrix is cast over a mold within a PDMS support and sandwiched between two coverslips to form two open channels. One channel is filled with densely-packed cells and plugged to create a tumor-like cell reservoir. White arrows indicate diffusion of nutrients from surrounding medium to cell reservoir. Magnified view shows invasion from cell reservoir to open channel.

We fabricated a simple device to mimic topographical features of the perivascular niche **(Fig 1C)**. The device consisted of a tumor-like cell reservoir adjacent to a parallel open channel, which were both embedded in 3D matrix. To form the device, we first fabricated two spacers to control the distance between channels and vertical distance at which the channels were suspended. First, we fabricated a horizontal spacer to control the distance between the parallel channels **(Fig 2A).** Vertical spacers consisted of a thin layer of PDMS. We then combined horizontal and vertical spacers to mold a PDMS holder with parallel, suspended channels **(Fig 2B,3A)**, which was then sandwiched between two glass coverslips which were fastened together **(Fig 3B)**. The PDMS holder was used to support wires as a mold upon which hydrogel was cast **(Fig 3C)**. Removal of the wires resulted in two ~170 µm diameter parallel channels embedded within ~500 µm thick 3D matrix. We fabricated gels with a shear modulus of ~300 Pa. This modulus is both within the range of the values typically reported for brain tissue (300 – 3000 Pa)^25,26^ and conducive to maintaining channel integrity during fabrication. One channel was seeded with cells at high density and then plugged on each end to restrict cell migration to the hydrogel only **(Fig 3D)**. The entire device was then bathed in cell medium, and the other channel was left open to allow passive filling. The cell reservoir and vessel mimic were separated by ~500 µm of 3D matrix through which cells invaded to reach the open channel. Nutrients from medium could diffuse to the cell reservoir either through the open channel or at the sides of the device through the 3D hydrogel.

**Figure 2.**
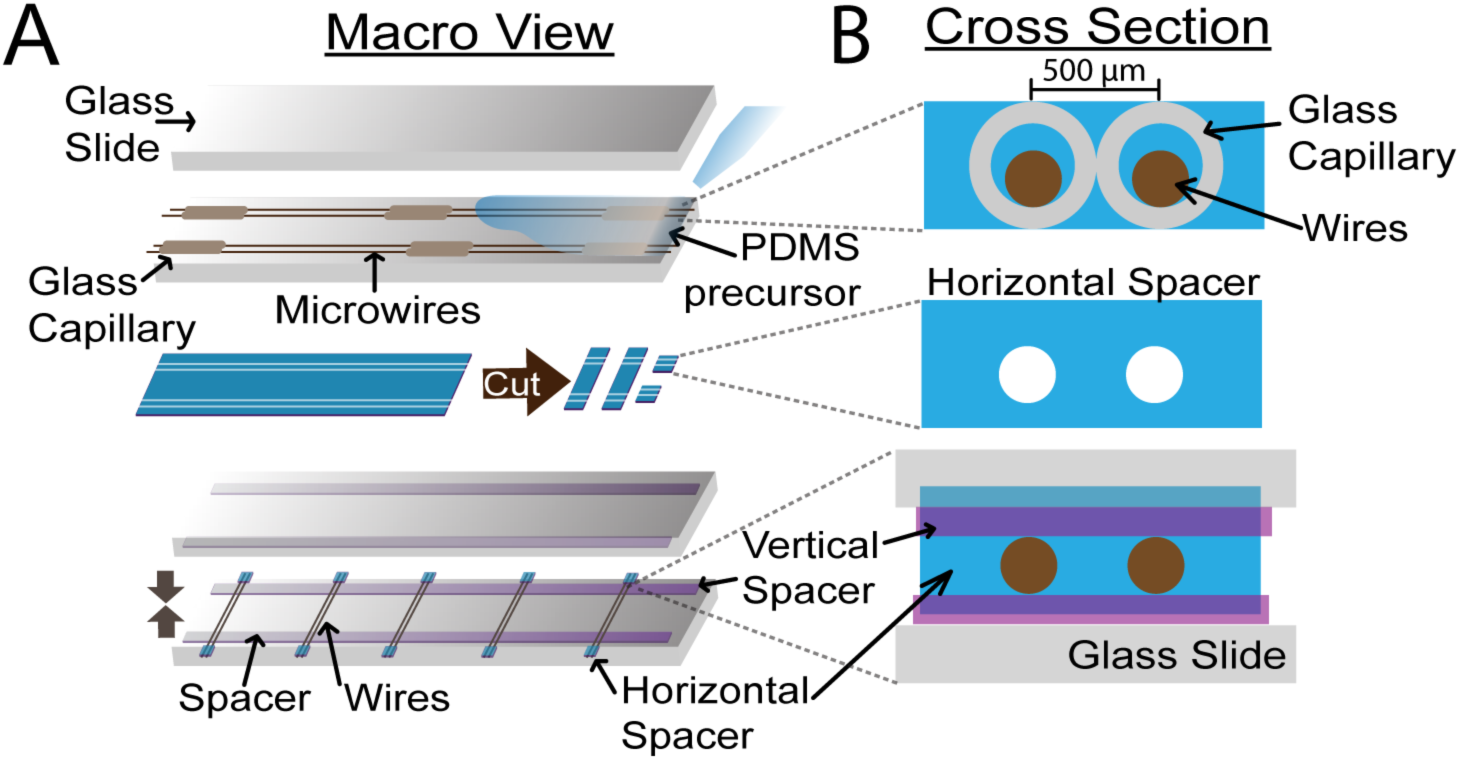
Spacer fabrication to align parallel channels suspended in PDMS. Glass capillaries of known diameter are used to control spacing between microwires during horizontal spacer fabrication. Horizontal spacers are cut from PDMS and then used to control wire-to-wire spacing during PDMS support fabrication. Vertical spacers are used to suspend microwires A) Macro view showing glass slide and wire alignment. B) Close-up view of cross section of spacer alignment showing control over vertical and horizontal dimensions.

**Figure 3.**
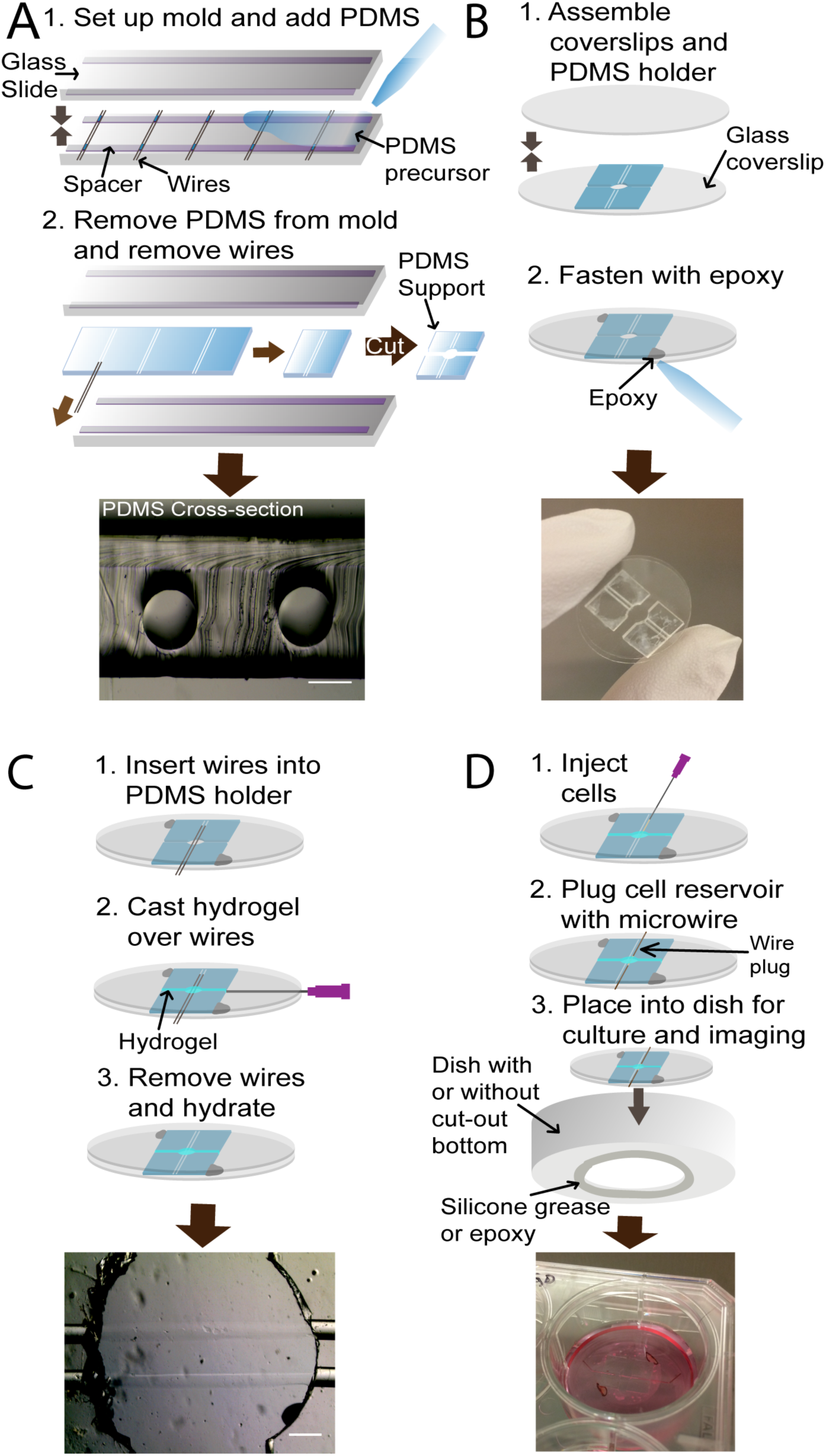
Device fabrication. A) PDMS supports are fabricated by casting PDMS over a wire mold, curing, and cutting the mold. Scale = 200 µm. B) PDMS supports are placed between coverslips and fastened with epoxy. C) Wires are inserted into the device to mold hydrogel, and matrix is cast over the wires. Wires are removed leaving open channels. Scale = 500 µm. D) Cells are injected into the reservoir channel, and plugs are added to prevent cells from migrating out of channel.

### 3D Topographical Model Promotes Tumor Expansion, Invasion in Masses, and Invasion along the Channel

To test whether GBM cells would indeed undergo multi-step invasion and follow topographical features within the matrix, we seeded U87-MG human GBM cells into seven devices and tracked invasion for 2-3 weeks. Based on our previous studies, we selected 3 wt% HA matrix functionalized with 0.5 mmol/L RGD as our matrix^27,28^ and then tracked tumor expansion and invasion for 2-3 weeks. We chose a 3wt% HA concentration to facilitate fabrication of gels with the desired shear modulus of 300 Pa with low enough matrix density to enable cell invasion. We have previously observed in 2D studies that an RGD concentration of 0.5 mmol/L promotes cell spreading and formation of broad lamellopodia.^28^ Cells began to invade the matrix approximately 4 days after seeding **(Fig 4A**) and continued to invade in multicellular masses over the next several days, tunneling through the matrix toward the open channel. Cells began to reach the channel around day 13 and reoriented to follow the channel wall **(Fig 4A)**. Within the channel, cells generally adopted a linear morphology and were spaced further apart than cells packed densely in invading masses.

**Figure 4.**
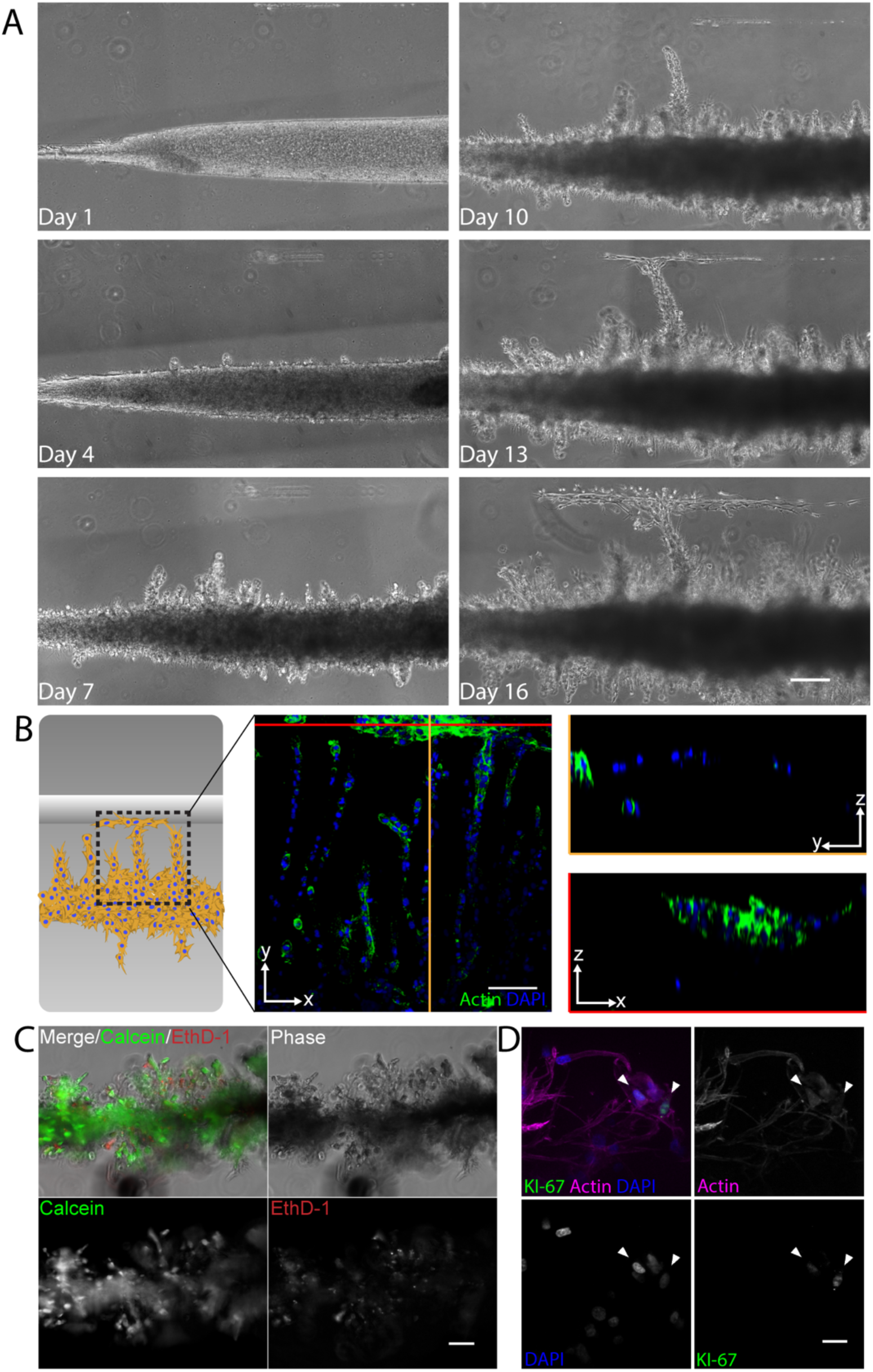
Tumor expansion, 3D invasion, and track-based invasion within the PVN model. A) Time series for cell expansion and invasion in HA-RGD. Scale = 200 µm. B) Schematic of region of interest within gel for z-stack (240 µm, 5 µm slices) demonstrating that cells generally migrate in the *xy* plane and that the cells begin to follow the channel after entering. Slice in *xz* plane shows migration along the *x* axis within the channel and slice in *yz* plane shows minimal migration in the *z* direction. Scale = 100 µm. C) Calcein AM to stain viable cells and EthD-1 to stain apoptotic cells demonstrates cell viability within the tumor reservoir. Scale = 100 µm. D) Z-stack (56 µm in height) of invading cells demonstrating relatively few Ki-67 positive nuclei (indicated by arrowheads). Scale = 20 µm.

Cells invading form the reservoir to the channel were primarily coplanar in *xy*, tunneling directly toward the open channel and then migrating along the channel wall (**Fig 4B).** This highly anisotropic migration suggests the presence of strong, diffusive chemotactic gradients between the reservoir and channel. While the lack of migration in *z* may initially seem surprising, solute diffusion is strongly promoted in the *xy* plane due to the device being covered on top and bottom by coverslips but being open to the medium in all lateral aspects (*xy* plane). Passive filling of the open channel with medium likely facilitates steeper chemotactic gradients of some medium components due to close proximity to the cell reservoir, further inducing cells to invade directly toward the channel.

To determine whether cells remained viable within the tumor reservoir, we performed a cell viability assay after 16 days in culture and found that most cells stained positive for calcein AM, but not ethidium homodimer (EthD-1) (**Fig 4C**). Cells that did stain positive for EthD-1 were not localized to any particular region of the tumor reservoir. We did not observe a necrotic core, which has been observed in larger tumorsphere models (>500 µm diameter).^45^ Rather, the cells in the core of the 170 µm diameter reservoirs remain viable despite the high cell density. This is consistent with our observation that the majority of cells within the tumor reservoir are mobile during live imaging experiments. Despite high cell viability, we observed relatively few Ki-67 positive cell nuclei (**Fig 4D**). Cells with nuclei positive for Ki-67 did not appear to be localized to any particular region of the device; however, more rigorous testing is required to conclusively determine whether any particular patterns exist. While a number of variables in culture conditions could be responsible, the observed low number of proliferating cells may support the “go or grow” hypothesis. This hypothesis suggests cells upregulate either invasive mechanisms or proliferative mechanisms, but not both.^46,47^ Our platform should be useful for future investigation of the effects of topographical microenvironment, matrix composition, and culture or drug-treatment on proliferative and invasive phenotypes.

We next sought to observe both expansion and invasion within the model. Invasion in the HA-RGD matrix was visually distinct from expansion of the reservoir mass in phase contrast appearing as a dark region of high cell density (**Fig 5A).** Because invasion occurred mostly in the *xy* plane, we used the area as an approximate measure of expansion and invasion **(Fig 5A).** We tracked reservoir growth and cell invasion for 16 days until the first device was fixed for further analysis, normalizing the areas to the total area on day 1 to account for any variance in cell seeding and channel length.

**Figure 5.**
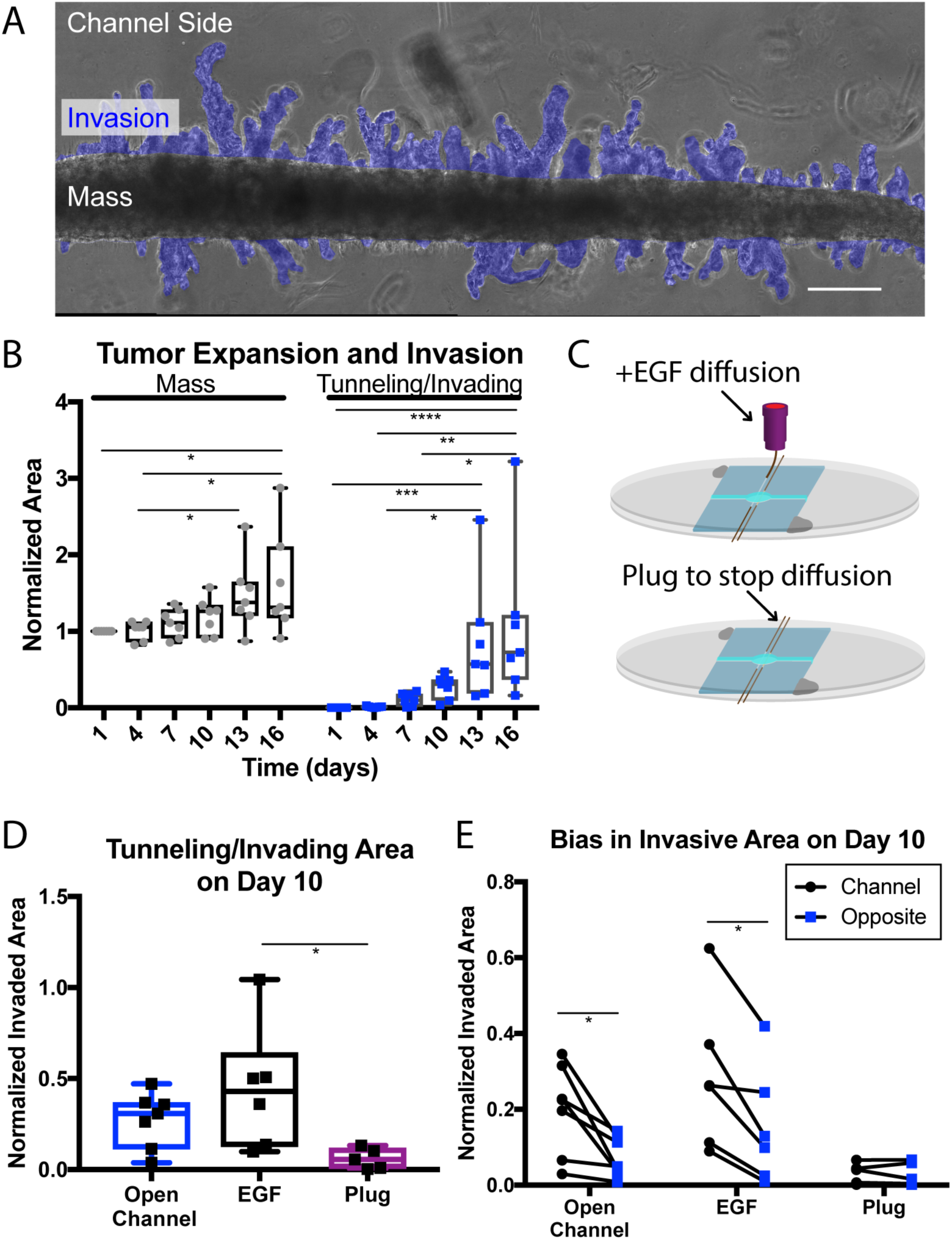
Cell expansion and invasion. A) Example of invasion quantification showing area of invasion (blue) as compared to the center mass of cells. Scale = 200 µm. B) Mass area and tunneling/invading area normalized to the initial area for n=7 devices over the first 16 days after seeding. *, p<0.05; **, p<0.01; ***, p<0.001; ****,p<0.0001 by Friedman’s Test followed by Dunn’s test for multiple comparisons. Center line represents median, boxes represent 25^th^ and 75^th^ percentiles, whiskers represent min and max. C) Device schematic for EGF diffusion into the channel and plugged channel to prevent diffusion. D) Total area of invading and tunneling cells is higher in the EGF condition compared to plugged channel condition. *, p<0.05 by ANOVA followed by Tukey’s test for multiple comparisons, boxes represent 25^th^ and 75^th^ percentiles, whiskers represent min and max. E) Invading area (blue) on the side nearest the channel compared to the opposite side on day 10 normalized to the total initial area for each device. Each of the lines represents results obtained with a different device. Significantly more cellular expansion and infiltration is observed on the channel side in devices containing an open channel and EGF diffusion within the channel, but not with a plugged channel (n=5-7). *, p<0.05 by paired t-test of average difference between channel side and opposite.

Both the reservoir area and the area of invasion grew over the 16 days **(Fig 5B)**. The reservoir area began to expand earlier than the invaded area in most devices, beginning around day 4 and expanding more rapidly around day 13. The area of invasion did not begin to increase until around day 7, but then expanded more rapidly than the reservoir area. By day 16, the increases in area were due largely to invasion as opposed to reservoir expansion. These data suggest that some threshold, possibly time or a certain degree of confinement, is required for cells to upregulate invasive mechanisms. Once invasion begins, it proceeds more rapidly than reservoir expansion.

We also observed that invasion was generally biased toward the open channel, despite the fact that the cell mass was surrounded on all sides by an HA matrix bathed in culture medium. We hypothesized that this may be due to a chemotactic effect arising from more rapid diffusion of serum medium into the open channel. The cell reservoir was much closer to the open channel (~500 µm) than to the gel-bath interface at the sides of the device (~4 mm). The open channel in close proximity to the reservoir may allow for increased medium diffusion and serve as a chemoattractant. To further investigate the effects of chemotactic gradients within the matrix, we set up two additional conditions. First, we sequestered EGF-containing medium within the channel (**Fig 5C**) to allow for EGF diffusion into the channel but not into the surrounding medium. We also set up a control experiment in which we used another wire to plug the open channel on each end, preventing medium diffusion into the open channel (**Fig 5C**). First, we quantified the total area of cell invasion after ten days and found that cells invaded significantly more area in devices treated with EGF compared to those with plugged channels (**Fig 5D**). We then quantified and compared the invasion area on the side of the reservoir nearest the open channel with the invasion area opposite. In both the open channel condition and the EGF perfusion condition, we observed significantly more invasion toward the parallel channel compared to the opposite side (**Fig 5E**). Together, these results suggest that serum can induce both chemotaxis and chemokinesis, with EGF further enhancing chemokinesis. This also demonstrates that the PVN model can serve as a platform for investigating the effects of other soluble factors within the context of matrix and topographical features.

### Cell Morphologies in the PVN Model Resemble Tumor Cell Invasion Toward and Along Vascular Tracks

Cell morphology is one indicator of whether cells are interacting with the microenvironment in a manner that captures important features of GBM. We investigated cell morphology in different stages of invasion and locations in the gel (**Fig 6A**). Long, thin protrusions generally preceded 3D invasion (**Fig 6B**). These protrusions varied in thickness and length. While most protrusions extended between 10-50 µm, protrusions occasionally reached >100 µm in length (**Fig 6C**). The number, length, and thickness of the protrusions more closely resembled dendritic processes similar to those that have been observed *in vivo*.^8,21,35^ This cell morphology contrasts elongated cells exhibiting a few short protrusions that can be observed in fibrillar matrices.^48^ Together, these results suggest that the HA matrix supports cell invasion that is morphologically similar to that observed in brain parenchyma.

**Figure 6.**
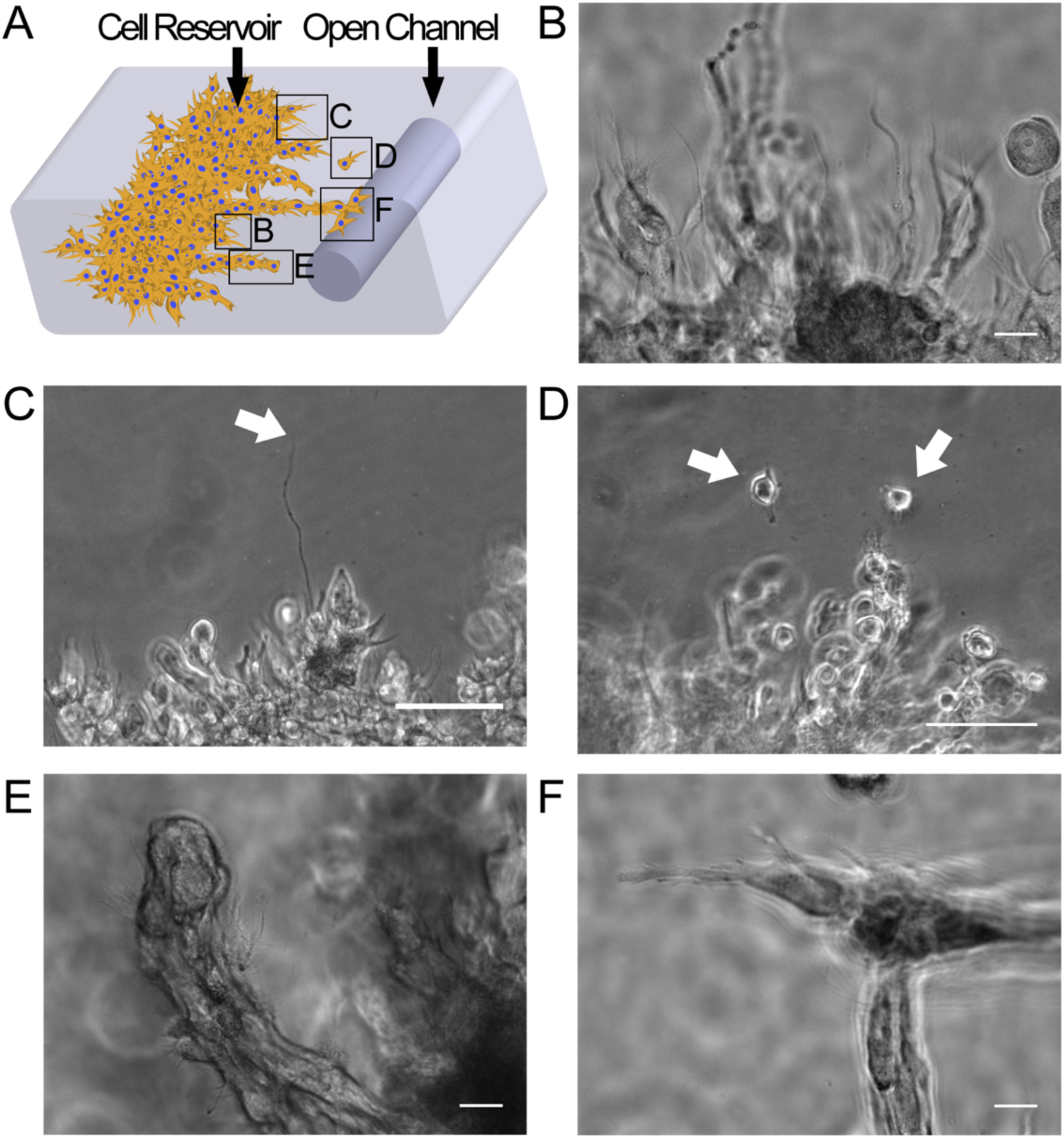
Cell morphologies during invasion. A) Schematic representing the variety of invasion morphologies depending on location and stage of invasion. Letters correspond to subsequent figure parts. B) Invasion is usually preceded by the extension of long protrusions. Scale = 20 µm. C) Protrusions occasionally reach lengths >100 µm, end of protrusion marked by arrow. Scale = 100 µm. D) Occasionally single cells are observed invading 3D matrix (arrows). Scale = 100 µm. E) Most cells invade in multicellular “tunnels”. Scale = 20 µm. F) Upon reaching the open channel, cells rapidly change directions and follow the channel wall. Scale = 20 µm.

We also investigated how cells move on a collective level. GBM cells *in vivo* move collectively as multicellular strands and less frequently as single cells.^8,35,48–50^ While single cells occasionally invaded 3D matrix (**Fig 6D**), invasion was most commonly observed as multicellular masses or “tunnels,” with protrusions extending from the end and sides of these masses (**Fig 6E**). This dynamic is consistent with a mechanism in which cells degrade and/or tunnel through the dense 3D matrix during migration. After reaching the open channel, cells changed direction and morphology (**Fig 6F)**. Cells in the open channel were less rounded and more elongated, aligning parallel to the length of the channel. Thus, an embedded, open channel serving as a topographical model of the PVN is sufficient to recapitulate changes in cell morphology during invasion.

### Topographical Transitions are Observed in Multiple 3D Matrix Types

One of the key features of GBM invasion *in vivo* that we aimed to recapitulate in this model is rapid migration of cells along vascular tracks relative to the slow interstitial invasion. Having established that cells change morphology and direction to follow the vessel-mimetic open channel, we investigated whether cells in channels migrated more rapidly than cells in the 3D matrix. Furthermore, we asked whether transitions were induced only by the HA matrix, known to promote cell invasion in the PVN.^51–53^ We thus chose collagen I, a fibrous matrix not normally found in brain ECM. Choosing a collagen matrix also enabled us to investigate the degree to which migration mode is dependent on matrix structure and to explore the versatility of our device in supporting the use of other matrix types.

We observed cell invasion through 3D matrix and channels in both HA-RGD and collagen I hydrogels and compared cell speeds in these two regions of interest for both matrix types (**Fig 7A**). Overall, cells in collagen I were more elongated with a more mesenchymal morphology and migrated more rapidly than cells in HA-RGD by approximately a factor of four. In contrast, cells in HA-RGD were more round with numerous dendritic protrusions. Despite the contrast in morphology and overall speed, cells invaded more rapidly (approximately twice the speed) in channels than in the 3D matrix for both matrix types. While the overall speed of invasion is dependent on matrix type, the topographical cues from the channel drive a change in cell speed that is independent of matrix composition.

**Figure 7.**
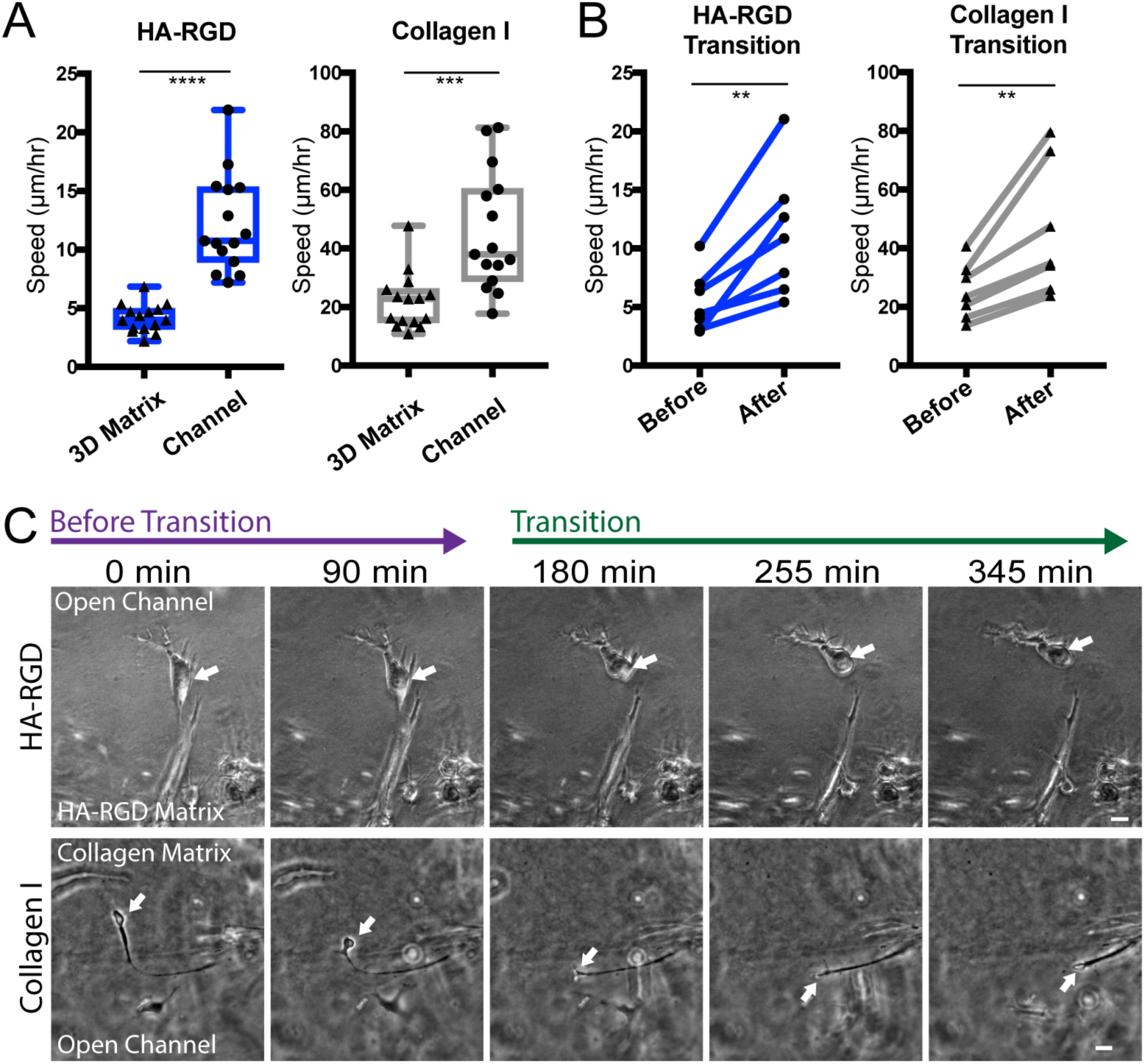
Transitions in direction, speed, and morphology as cells encounter the open channel. A) Cell speeds in 3D matrix are slower than in channels, ***, p<0.001 by student’s t-test. Center line represents median, boxes represent 25^th^ and 75^th^ percentiles, whiskers represent min and max. B) Single cell speeds increase after the transition, **, p<0.01 by paired t-test of average difference between speed before transition and speed after transition. C) In both HA-RGD and collagen I matrices, cells change direction to align with the channel. Arrows point to cell nucleus. Frames at 0 min and 90 min are before the transition of the entire nucleus out of the 3D matrix while subsequent frames occur after the transition. Scale = 20 µm.

### Topography-Driven Changes in Migration Speed are Instructive rather than Selective

The degree to which increased speed exhibited by cells within channels was due to cell-ECM interactions as opposed to cell-cell interactions remained unclear. For example, tumor cells often invade in a communal fashion, with leader cells remodeling matrix to enable rapid migration of follower cells.^54^ It is possible that the difference in migration speed in each topology is driven not by cell-instructive cues but rather by different subpopulations of cells, with one subset more adept at 3D invasion and another more adept at perivascular invasion. Furthermore, the cell-ECM and cell-cell interactions that underlie single and collective cell migration could depend on the matrix type. Cells in the collagen matrix were more frequently observed moving as single cells and did not invade in the multicellular masses as in HA-RGD. It is likely that the increased pore size and fibrous structure of collagen was more permissive of cell invasion, either changing or eliminating the role of leader cells.

To begin assessing the relative role of single cells in the transition to vasculature invasion, we quantified the invasion of cells crossing from 3D matrix to the open channel (**Fig 7B)**. A transition was defined as the time at which the nucleus fully exited the 3D matrix into the channel. We measured cell speed for a total of 6 hr with a transition occurring between the first 3 hr and the second 3 hr. Migration speeds before and after the transition were compared (**Fig 7B,C**). Again, cells in collagen I matrix migrated rapidly compared to HA-RGD. However, in both matrices, individual cells increased their speed after the transition. The speeds before and after the transition were similar to those measured when tracking cells in 3D matrix or the channel only. We concluded that single cells responded to the open channel topography by increasing speed, leading to overall more rapid invasion along the channel.

### Cytoskeletal Architecture Varies by Location and Matrix Type

While increases in cell speed were observed in both matrix types, phase imaging suggested vastly different cell morphologies (**Fig 7C**). Specifically, we observed that cells in collagen I or in channels tended to have more linear morphologies in contrast to the rounded, dendritic morphologies seen in HA-RGD. Cytoskeletal architecture, particularly that of the actin cytoskeleton, can reflect mechanisms underlying invasion.^48^ We asked whether distinct cytoskeletal architectures were correlated with specific invasion patterns seen in our various matrix formulations (HA, collagen I) and topographies (3D, channel). Following past studies, we reasoned that actin bundles and stress fibers are more likely to accompany the fast mesenchymal-like migration observed in the open channels and in collagen.^55–57^ In contrast, more diffuse cortical actin would be expected within HA-RGD. We therefore applied structured illumination microscopy (SIM) to characterize actin cytoskeletal architecture in each of these scenarios **(Fig 8**).

**Figure 8.**
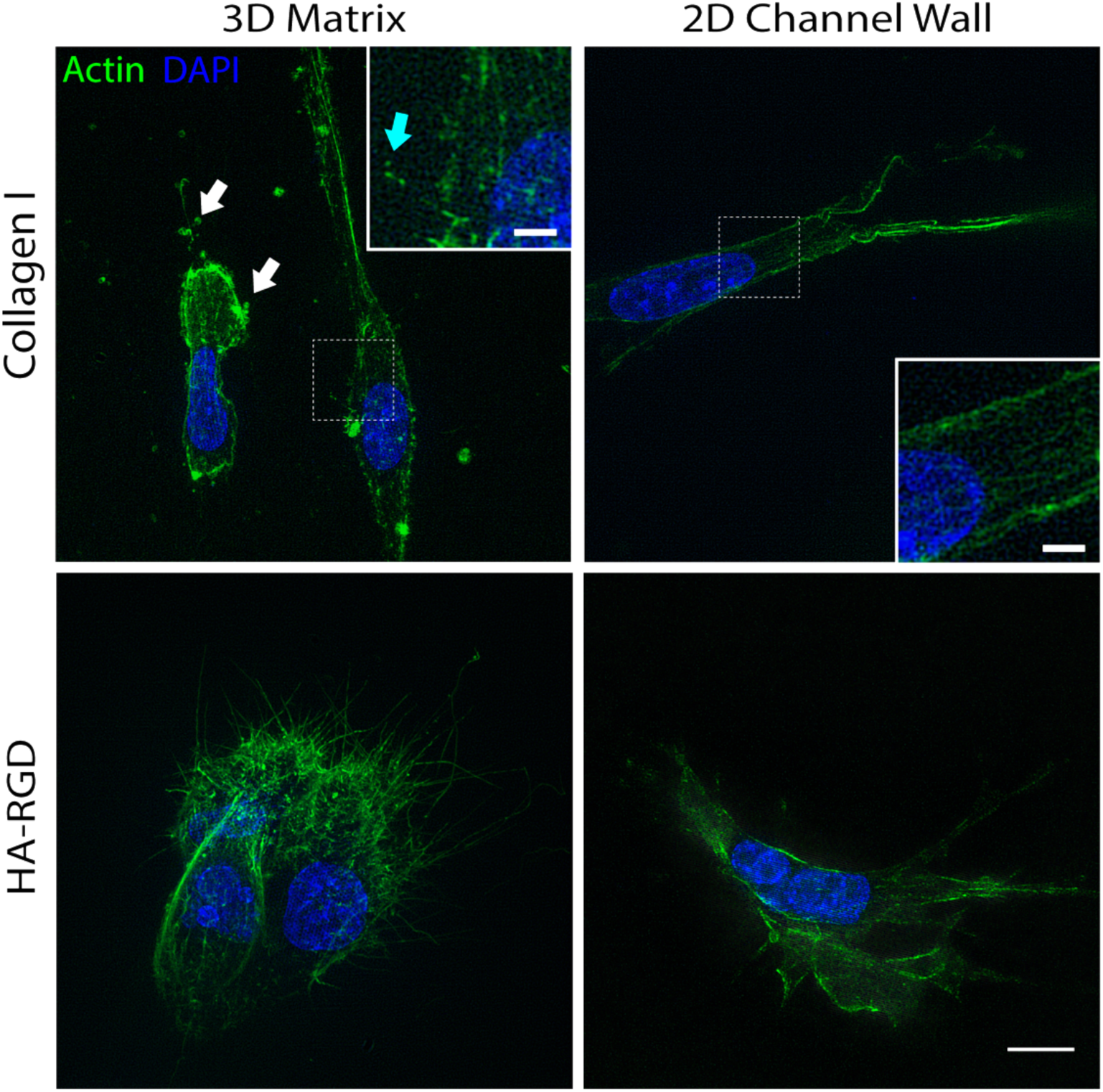
SIM imaging of fixed cells revealing distinct actin architectures. Cells in 3D collagen I gels show evidence of membrane blebbing (white arrows) and nuclear squeezing and also express more numerous short filopodia (blue arrow, inset) than cells in the collagen I channel. Cells in 3D HA-RGD matrix are densely packed, rounded, and express numerous long, thin filopodia. Cells in HA-RGD channels express actin filament bundles and protrusions aligning with channel wall. Scale = 10 µm, inset scale = 2 µm. Z stacks were 20 µm (top left), 11 µm (top right), 23 µm (bottom left, and 11 µm (bottom right) with 1 µm spacing between slices.

As we anticipated, cells in 3D collagen I assembled long actin filament bundles that align with the direction of migration. This was not the exclusive phenotype, as some cells exhibited an elongated nucleus with membrane blebs, suggesting confined, amoeboid migration. Compared to cells in collagen I channels, cells in 3D collagen expressed short and more numerous filopodia. In contrast to cells in collagen, cells in 3D HA were almost exclusively rounded and had numerous single actin filaments with relatively thin and few filament bundles. Cells in HA-RGD channels were elongated with fewer and longer actin filament-based invasive protrusions. As with collagen I, this was accompanied by assembly of actin bundles aligned in parallel with the channel. Overall, the morphology of cells in 3D were distinct from those in the channel, and the contrast was much greater in HA-RGD gels. These results explain the hierarchy of migration speeds (channel collagen > 3D collagen > channel HA > 3D HA). Migration speed is tied to the propensity of the matrix to support actin bundle-driven mesenchymal migration: fastest in a fibrous matrix arranged in a linear channel, and slowest in a non-fibrous matrix in 3D. These results also underscore that migration mode is dependent on matrix type, and that cell morphologies in HA-RGD matrix compared to collagen I are more similar to those observed in vivo. Still, topography influences migration mode and speed in a similar fashion within a particular matrix type. While we observed an analogous increase in speed and change in morphology within HA-RGD and collagen I matrices, it is possible that changing cell types or matrix compositions will result in an altogether different type of response to topographical cues. Future investigation of other matrix compositions, possibly with spatial organization of ligands or modulus, may uncover potentially synergistic roles of topography and matrix in promoting invasion.

### Integrin Engagement and Cell Contractility Promote Cell invasion in HA Matrices

We then investigated whether integrin engagement and cell contractility were necessary to support invasion into HA-based matrices. While cells are capable of binding to HA-based matrices independent of integrin adhesions, we hypothesized that integrin binding and contractility allow cells to squeeze through matrix while forming reinforced adhesions that facilitate cell invasion. We compared mass expansion and invasion areas in devices with or without RGD in the matrix and with or without myosin II inhibition through blebbistatin treatment. Blebbistatin treatment significantly reduced the mass expansion of the reservoir **(Fig 9A)**. Cells in devices lacking RGD or treated with blebbistatin exhibited significantly less invasion and tunneling into the surrounding matrix **(Fig 9B,C)**. Furthermore, we observed different cell morphologies in matrices with or without RGD. Cells invading HA-RGD exhibit long, thin protrusions preceding invasion while cells in HA matrix remain largely rounded (**Fig 9D**). Thus, cells appear to transition from a slow, more amoeboidal mode of migration with no RGD and low contractility to a more rapid, mesenchymal mode of migration with RGD in the matrix and uninhibited contractility.

**Figure 9.**
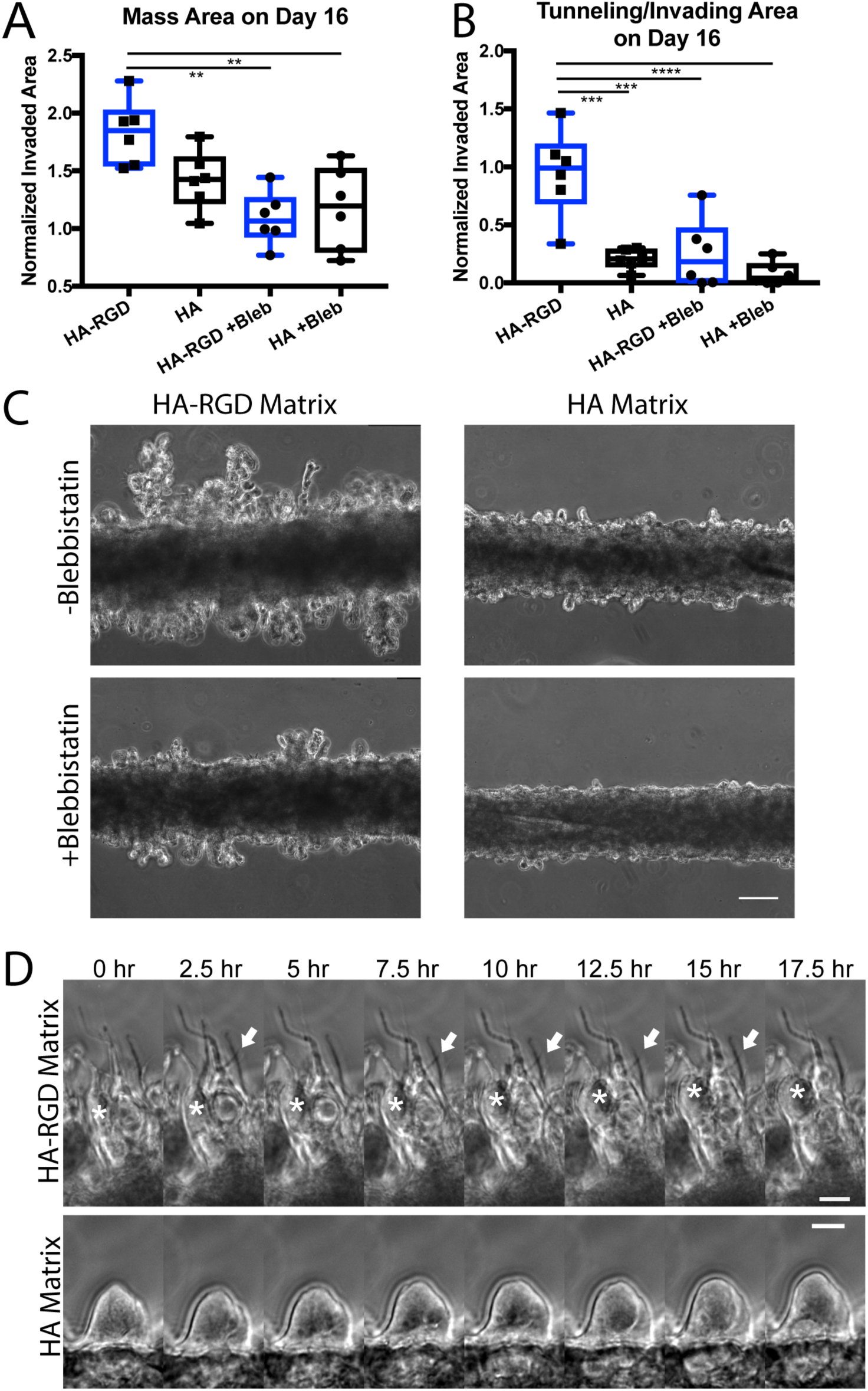
Cell invasion is dependent on integrin engagement and cell contractility. A) Mass area and B) tunneling/invading area of cells in in HA-RGD and HA matrices treated with or without 10 µM blebbistatin normalized to the initial area on day 1 for n=6 devices per condition 16 days after seeding. **, p<0.01; ***, p<0.001; ****, p<0.0001 by ANOVA followed by Tukey’s test for multiple comparisons. Center line represents median, boxes represent 25^th^ and 75^th^ percentiles, whiskers represent min and max. C) Tunneling and invading morphology is decreased with blebbistatin treatment and without RGD ligand in matrix. Scale = 100 µm. D) Invasion in HA-RGD matrix occurs is preceded by thin protrusion extension while invasion in HA matrix is not. Arrow indicates protrusion dynamically extending and retracting and asterisk indicates cell shifting toward protrusions.

Compared with our previous results in collagen I matrices, these data suggest that our matrices promote a range of migratory phenotypes with more mesenchymal behavior (collagen I > HA-RGD > HA). These results may also reflect the role of pore size in regulating invasion. We have previously demonstrated that our HA gels are nanoporous (average mesh size ~100-150 nm).^27^ Thus, cells are likely to rely on matrix degradation to invade into the gel resulting in more amoeboidal invasion. Collagen gels, however, have pore sizes on the scale of tens of microns that enable cell migration without degradation, resulting in faster invasion speeds.^58^ As a whole, these experiments demonstrate the possibility of using our platform to investigate the effects of matrix composition and treatment, which in the future could be leveraged to perform deeper mechanistic studies or conduct screening.

## Conclusions

We have developed a topographical culture model that is sufficient to recapitulate and observe multiple stages of invasion (expansion, matrix infiltration, and invasion along anatomical tracks) with a distinct transition in cell morphology and speed resembling GBM invasion toward and along the PVN. We have demonstrated that increased speed and change in direction are generalizable across matrix types, and that distinct actin architectures support these transitions. This work underscores the utility of incorporating topographical cues into 3D invasion models to study multiple modes of invasion relevant to clinical GBM progression. While this study is focused on topographical effects on speed and cell morphology, the platform could be used to conduct screening and investigate cell signaling and mechanistic pathways underlying invasion in the PVN.

## Acknowledgements

Confocal images were acquired at the CRL Molecular Imaging Center at UC Berkeley, supported by National Science Foundation grant DBI-1041078. We would like to thank Holly Aaron and Jen-Yi Lee for their assistance in training. Structured illumination microscopy was conducted at the UC Berkeley Biological Imaging Facility, which was supported in part by the National Institutes of Health S10 program under award number 1S10(D018136-01). The content is solely the responsibility of the authors and does not necessarily represent the official views of the National Institutes of Health. We would like to thank Dr. Steve Ruzin and Dr. Denise Schichnez for their technical assistance and training. Finally, the authors gratefully acknowledge financial support from the following sources: National Science Foundation (Graduate Research Fellowship to K.W.); National Institutes of Health (Ruth L. Kirschstein Predoctoral Individual National Research Service Award F31GM119329 to S.L., R21CA174573, R21EB025017, and R01GM122375 to S.K.); and the W.M. Keck Foundation (S.K.).

